# Diet induced insulin resistance is due to induction of PTEN expression

**DOI:** 10.1101/2025.03.25.645201

**Authors:** Radha Mukherjee, Priya Pancholi, Malvika Sharma, Hilla Solomon, Merna N. Timaul, Claire Thant, Rory McGriskin, Omar Hayatt, Vladimir Markov, John D’Allara, Simona Bekker, Jacqueline Candelier, Sebastian E. Carrasco, Elisa de Stanchina, Kiran Vanaja, Neal Rosen

**Affiliations:** Program in Molecular Pharmacology, Memorial Sloan Kettering Cancer Center; New York City, NY 10065, USA; Roux Institute, Bioengineering Dept., Northeastern University; Boston, MA 02120; Laboratory of Comparative Pathology, Weill Cornell Medicine, Memorial Sloan Kettering Cancer Center, and Rockefeller University; New York City, NY, 10065, USA; Department of Pathology and Laboratory Medicine at Weill Cornell Medicine; New York City, NY 10065, USA

## Abstract

Insulin resistance is a condition associated with obesity, type 2 diabetes(T2D), hyperinsulinemia, hyperglycemia and defined by reduced sensitivity to insulin signaling. Molecular causes and early signaling events underlying insulin resistance are not well understood. Here we show that insulin activation of PI3K/AKT/mTOR signaling in insulin target tissues, causes mTORC1 induction of PTEN translation, a negative regulator of PI3K signaling. We hypothesized that insulin resistance is due to insulin dependent induction of PTEN that prevents further increases in PI3K signaling. In a diet induced animal model of obesity and insulin resistance, we show that PTEN levels are increased in fat, muscle, and liver. Hyperinsulinemia and PTEN induction are followed by hyperglycemia, severe glucose intolerance, and hepatic steatosis. In response to chronic hyperinsulinemia, PTEN remains increased, while AKT activity is induced transiently before settling down to a PTEN-high and AKT-low state in the tissues, predicted by computational modeling of the PTEN-AKT feedback loop. Treatment with PTEN and mTORC1 inhibitors prevent and reverse the effect of PTEN induction, rescue insulin resistance and increase PI3K/AKT signaling. Thus, we show that PTEN induction by increased insulin levels elevates feedback inhibition of the pathway causing insulin resistance, its associated phenotypes, and is a potential therapeutic target.

## Main Text

An increase in sedentary lifestyle and the dietary consumption of carbohydrates and fats leads to multiple metabolic diseases such as insulin resistance, obesity, and Type 2 Diabetes (T2D). T2D is a condition that is defined by insulin resistance, hyperglycemia, and hyperinsulinemia unless it occurs with beta cell failure and insulin deficiency(*1*). Insulin resistance and pre-diabetes precedes T2D and is also a driver of multiple other metabolic syndromes such as metabolic dysfunction associated steatohepatitis (MASH) and atherosclerosis(*2–4*). Insulin exerts its effects by binding to and activating insulin receptor (IR) signaling(*5–10*). Insulin resistance is defined by a requirement for increasing concentrations of insulin to activate IR-PI3K-AKT signaling in peripheral tissues like muscle, adipose and liver. This results in insufficient uptake of glucose into these tissues and increased export of glucose from the liver(*11–14*). Increased caloric intake results in hyperinsulinemia that initially maintains normoglycemia but, with time, the rising levels of insulin cannot keep up and hyperglycemia ensues. Chronic hyperglycemia and hyperinsulinemia eventually results in islet cell exhaustion and apoptosis and insulin deficiency(*14*).

Insulin resistance is characterized by the reduction in IR signaling(*11–14*). Insulin binding to IR causes its activation and phosphorylation of insulin receptor substrates (eg. IRS1 and 2) which bind to and activate PI3 kinase and other effector proteins. PI3kinase phosphorylates its substrate PI(4,5)diphosphate (PIP2) leading to the production of the phosphoinositide PI(3,4,5) triphosphate(PIP3)(*5–9*). Accumulation of PIP3 leads to the activation of AKT kinases and other substrates(*9, 15*). AKT activation drives glucose uptake and glycogen synthesis by phosphorylating its substrates AS160 and glycogen synthase, inhibiting the former and activating the latter. Phosphorylation of AS160 causes the activation of vesicular transport of GLUT4 glucose transporter, allowing the translocation of the latter to the cell membrane and enhanced uptake of glucose(*16*). In the liver, AKT activation suppresses hepatic glucose production by inhibiting gluconeogenesis and glycogenolysis and stimulating glycogen synthesis(*16*). It also inhibits the transcription factors required for gluconeogenesis by phosphorylating and excluding the FOXO transcription factors from the nucleus. In adipose tissue AKT stimulates glucose uptake and inhibits lipolysis. This reduces the production of non-esterified fatty acids thereby further reducing hepatic glucose production(*16*). The importance of both PI3K and AKT activation in maintaining glucose homeostasis is demonstrated by the rapid and marked induction of hyperglycemia in humans by drugs that inhibit either enzyme. One of the major downstream effects of AKT is induction of TOR kinase activity, which in turn, induces cap-dependent translation, ribosomal biogenesis, lipid synthesis and other processes required for cell growth (*17–24*).

The output of the pathway is regulated by multiple AKT and mTOR dependent inhibitory feedback loops that limit the amplitude and duration of the signal and maintains pathway homeostasis. PTEN is a lipid and protein phosphatase that de-phosphorylates PIP3 to generate PIP2 thereby antagonizing PI3K activity(*25*). Recently we discovered that PTEN is regulated by the PI3K/mTORC1/4E-BP1 axis by cap-dependent translation in a negative feedback loop. This mechanism is a key determinant of pathway homeostasis(*26*). Oncogene and growth factor dependent activation of PI3K in cancer cells increased PTEN levels, thereby reducing PI3K/AKT signaling. By contrast, nutrient starvation leads to decreased PTEN expression and AKT re-activation(*26*).

A cohort of T2D patients from Japan were found to harbor polymorphisms in the 5’UTR region of PTEN that led to an increase in PTEN levels by increased protein translation. Expression of the mutant PTEN reduced insulin dependent AKT activation(*27*). In addition, patients with Cowden’s Syndrome and monogenic PTEN mutations that lead to its haploinsufficiency are more sensitive to insulin action and protected from insulin resistance(*28, 29*).This led us to hypothesize that the induction of insulin levels in people on a high caloric diet increases PTEN levels and by that mechanism, reduces AKT activation and decreases sensitivity of the cell to insulin stimulation.

### Western Diet in mice and insulin stimulation in cells increase PTEN expression

We have shown previously that inhibition of the PI3K network using selective inhibitors reduces PTEN levels driven by mTOR/4E-BP1 mediated cap dependent translation of PTEN (Fig 1A)(*26*). This reduction in PTEN leads to pathway reactivation, reducing the efficacy of PI3K inhibitors thus serving as a fundamental mechanism of self-regulation of the network(*26*). However, we wanted to investigate if PTEN increases with insulin stimulation in insulin sensitive tissues like fat, muscle, and liver. In addition, we wanted to know if chronic hyperinsulinemia leads to increase in PTEN and its consequences on insulin sensitivity, both *in vitro* and *in vivo*.

**Figure 1.**
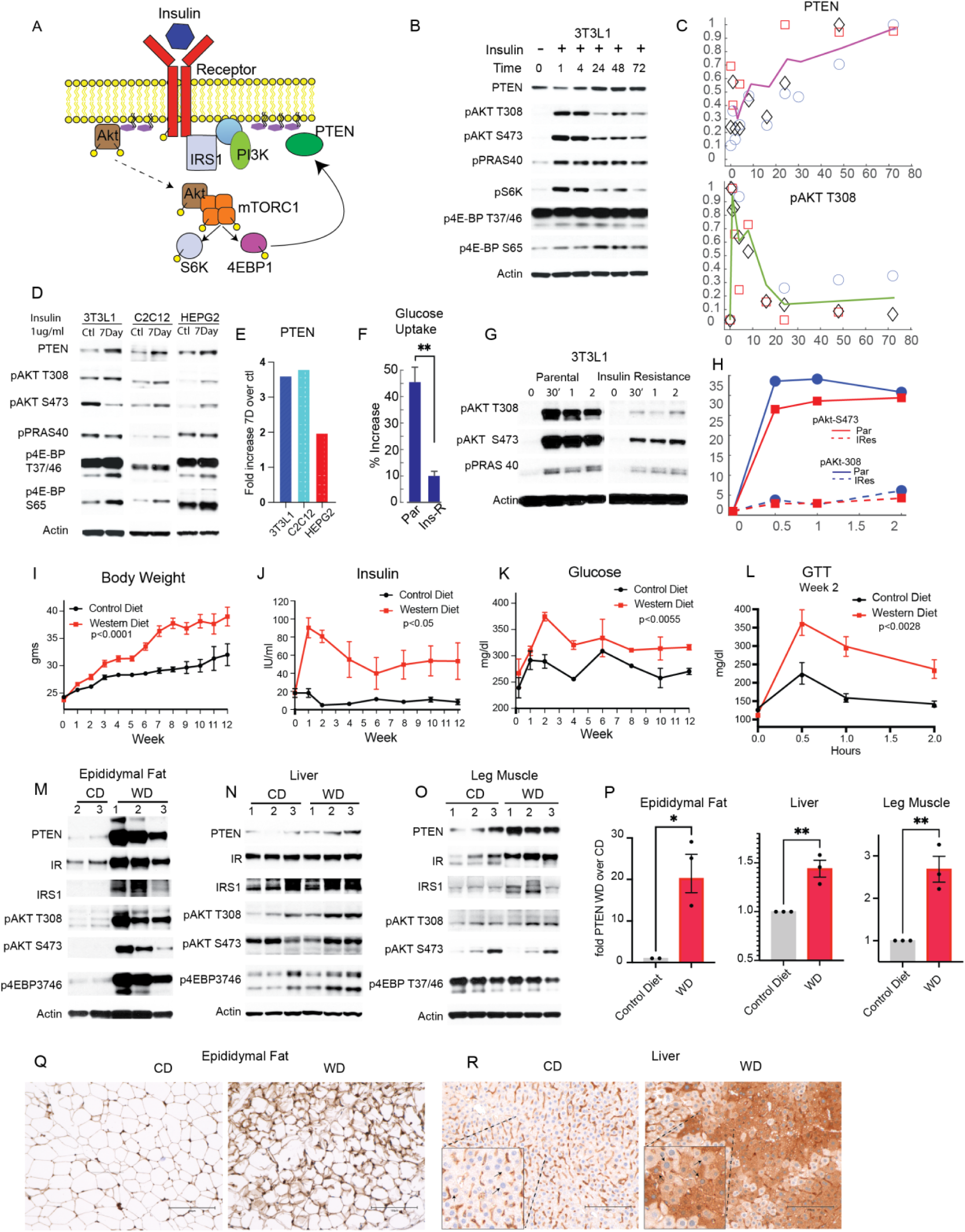
Western Diet in mice and insulin stimulation in cells increase PTEN expression. **(A)** Insulin/PI3K pathway diagram illustrating the PTEN feedback loop. **(B)** 3T3L1 cells were treated with 1ug/ml of insulin for the indicated times and analyzed for PTEN and PI3K pathway activity. **(C)** Quantification of PTEN and pAKT T308 from B and 2 other independent experiments. Mean and SEM plotted over time. **(D)** 3T3L1, HEPG2, C2C12 cells were treated with 1ug/ml insulin for 7 days and immunoblotted for the indicated targets. **(E)** Quantification of PTEN normalized to actin from D. **(F)** Glucose uptake assay in 3T3L1 cells after 7 days of insulin (Ins-R) or vehicle treatment (Par) and mean and SEM represented, p<0.01 calculated using student’s t test. **(G)** Immunoblotting of Par or Ins-R cells washed out and re-treated with 100ng/ml of insulin. H. Quantification of pAKT normalized to time t=0 from G. **(I-K)** C57BL/6J mice were either fed with regular/control (CD) or western diet (WD) for 12 weeks and at every time point food withdrawn for 3 hours and the weight, p<0.0001(n=10-15), insulin p<0.05 (n=5), glucose levels measured p<0.0055 (n=5). **(L)** After 2 weeks of control or western diet mice were fasted overnight and dosed with a bolus of glucose and their glucose levels measured for 2 hours at the indicated times for glucose tolerance test (GTT) p<0.01 (n=5). (I-L) Mean and SEM of n represented, and p values were calculated by a two-way ANOVA and post Bonferroni tests. **(M-O)** Mice were either fed with control (CD) or western diet (WD), food withdrawn for 3 hours and proteins extracted from epididymal fat (eWAT), liver and muscle and analyzed by western blotting (n=3, one of two independent experiments, also see Supplemental Fig S1N, protein from the eWAT from one control diet mouse could not be extracted). **(P)** Quantification of PTEN (normalized to actin) from (M-O) Mean and SEM of n plotted, **p<0.01, *p<0.05 calculated with unpaired t test with Welch’s correction. **(Q-R)** Epididymal fat (eWAT) and liver tissues from mice fed on CD or WD were immunostained for PTEN and representative images shown, (inset; indicated zoomed in area from R) (n=3, one of two independent experiments represented, also see Supp Fig 1SO).

We asked if insulin stimulation led to an increase in PTEN in fat, muscle, and liver cell lines. We found that when 3T3L1 adipose cells were treated with 1ug/ml of insulin, PTEN levels increased within 24 hours by 30% and remained increased until 72 hours by 60% and this coincided with the decreased kinetics of pAKT after its initial peak (Fig 1B-C). 4E-BP is a substrate of mTORC1 and a negative regulator of cap dependent translation(*30, 31*). Activation of mTOR and its phosphorylation of 4E-BP leads to its inhibition and thereby induces cap-dependent translation. In correlation with PTEN being high at late time points, phosphorylated 4E-BP remained increased upon insulin stimulation. In liver cell line HEPG2 and muscle myoblast cell line C2C12, PTEN increased upon insulin stimulation after 24 hours and remained increased for 3 days (Supp Fig 1A-B). Similar results were observed in cancer and normal cell lines upon stimulation with a cocktail of growth factors (Supp 1C-E)(*26*).

We next asked if continuous stimulation of fat, liver, and muscle cell lines with insulin, would lead to increased PTEN levels and reduced pathway sensitivity. We stimulated 3T3L1, C2C12 and HEPG2 with 1ug/ml of insulin for 7 days, replacing the media with insulin every day, and found PTEN to be increased by 3.5, 3.7 and 2 fold respectively in the insulin stimulated cells (Fig 1D). Phosphorylated AKT remained slightly increased, and its activity as measured by pPRAS40 either the same or reduced while p4E-BP remained increased in three all cell lines in correlation with increased PTEN levels. We investigated if the 7 day insulin treated cells had reduced responsiveness to insulin as compared to the insulin naïve cells. We stimulated the 3T3L1, HEPG2 and C2C12 cells for 7 days with 1ug/ml of insulin and then washed out the insulin and re-treated the cells with 100ng/ml of insulin. While the parental cells showed an increase of 25 and 35 fold in pAKT T308 and S473 respectively within 30 min of stimulation, the 7day insulin treated cells only increased their pAKT by 3-4 fold (Fig 1G-H), suggesting a severe decrease in insulin sensitivity in the 7 day insulin treated cells. The same phenomenon was observed in C2C12 and HEPG2 cells (Supp Fig F-G). Furthermore, glucose uptake in the 7 day insulin treated 3T3L1 cells was reduced to 10% as compared to 45% in the insulin naïve cells (Fig 1F). Therefore, PTEN increases during chronic prolonged insulin stimulation of adipose, liver and muscle cell lines and is associated with decreased insulin sensitivity of the cells.

We next hypothesized that a high fat high carbohydrate diet that causes hyperinsulinemia will also increase PTEN expression in insulin sensitive tissues in animals. We utilized a mouse model of diet induced insulin resistance and obesity(*32–34*) in which C57BL/6J 6 week old male mice were fed *ad libitum* with a “western diet” in which 41% of the calories are from fat and 43% from carbohydrates or control normal diet. The former simulates the modern dietary pattern in western countries characterized by high intake of processed foods rich in refined sugars, oils, and saturated fats. This diet when given to rodents mimics a variety of human metabolic syndromes including insulin resistance, obesity and liver steatosis(*35–37*).

Mice were fed on western diet (WD) and control diet (CD) and weight, insulin and glucose were measured over 12 weeks. Food was withdrawn for 3 hours before sampling at each time point since PTEN levels are affected by nutrient starvation over longer durations(*26*). Mice on the western diet and control diet started gaining weight within a week and, the former, gained significantly more weight than mice on CD (65% increase in WD and 31% increase in CD after 12 weeks) (Fig 1I). After 1 week, the serum insulin levels in mice on the western diet increased by 5-fold over that in mice on the control diet and remained increased for 12 weeks (WD over CD) (Fig 1J, Supp Fig 1H). Mice on the western diet showed increase in glucose levels 20% over those in control mice, 2 weeks after its initiation, and remained mildly hyperglycemic over 12 weeks (Fig 1K). Insulin sensitivity was measured using the glucose tolerance test (GTT) and a reduction in glucose tolerance began between 2 days and 1 week after western diet consumption and glucose tolerance was severely compromised by 2 weeks, coinciding with the start of hyperglycemia (Fig 1L, Supp Fig 1I). Glucose intolerance increased steadily over 6 weeks of the western diet (Supp Fig 1I) as demonstrated by the increase in peak glucose (30min after glucose stimulation) and adapted glucose levels (2 hours after glucose stimulation) of the GTT over 6 weeks, which remained increased by 55% and 46% respectively in mice on western diet over normal diet at the end of week 6 (Supp Fig 1J-K). As previously reported, we found that mice on the western diet developed hepatic lipidosis and steatosis (Supp Fig 1L) and a significant increase in leptin levels (11-fold over mice on CD) within 4 weeks (Supp Fig 1M)(*2, 4*). Altogether these data indicate that upon exposure to high fat diet, these mice developed insulin resistance and the onset of related metabolic pathologies.

We asked whether PTEN expression was increased in insulin sensitive tissues (epididymal white adipose tissue (eWAT), muscle, and liver). PTEN expression was significantly increased in eWAT and liver tissues after 1 week of western diet and in muscle after 2 weeks (18-fold in eWAT, 1.5-fold in liver and 2.7-fold muscle) as demonstrated by immunoblotting and immunohistochemistry (Fig 1M-P, Q-R Supp Fig 1N-O). The increase in PTEN expression measured by immunohistochemistry in the liver was very striking where hepatocytes (inset; in black arrows) showed a range of increased PTEN expression as compared to the mice on control diet (Fig 1R, Supp Fig 1O). Phosphorylated AKT T308 and S473 also increased in eWAT and AKT T308 phosphorylation was slightly increased in liver and was the same in the muscle while AKT S473 phosphorylation was unchanged in both liver and muscle (Fig 1M-O, Supp Fig 1N). Thus, in mice on the western diet, insulin activates the expression of PTEN, a negative regulator of upstream elements of the pathway. IRS1 and IR are other well-established negative feedback of the insulin receptor network which have been shown to decrease with insulin stimulation^6^. We found levels of insulin receptor and IRS1 increased in eWAT and were unchanged in the other two tissues (Fig 1M-O, Supp Fig 2A). In eWAT at 1 week, elevated insulin levels activate AKT T308 significantly, enough to maintain normoglycemia initially (Fig 1M, K).

These data show that PTEN increases in both *in vitro* and *in vivo* models of insulin resistance. In animals on a Western diet, increase in weight, serum insulin levels, and insulin resistance is accompanied by increase in PTEN expression within a week in insulin target organs.

### Chronic western diet increases PTEN, decreases AKT activity and is predicted by a computational model

By virtue of PTEN being connected to AKT in a feedback loop(*38*), it is expected that an increase in insulin and AKT signaling will result in PTEN increase. This increase in PTEN is less instantaneous and more integrative as a function of the amplitude and duration of AKT signaling. This may continue until a point where the increasing PTEN because of its inhibitory role will begin to cause smaller increases in AKT signaling. Over time this may then consequently reduce PTEN which will then in-turn cause slightly increased AKT signaling. This cycle will continue resulting in oscillations in the observed value of PTEN and AKT ultimately resulting in the establishment of a higher steady state PTEN and much reduced AKT signaling than the animals on control diet.

We investigated changes in PTEN and PI3K/AKT activity over chronic feeding of the western diet. Epididymal white adipose (eWAT), muscle and liver tissues were collected from mice that were on western or control diet as a function of time over 10 to 12 weeks (as mentioned earlier, food was withdrawn for 3 hours before sampling at each time point). In eWAT, we found PTEN to be significantly increased at 6, 8 and 10 weeks by 2 to 3-fold in mice on western diet over mice on control diet (Fig 2A-B). AKT phosphorylation on the other hand was significantly higher at 1 week (Fig 2C, Supp Fig S2C) in mice on western diet but by weeks 8 and 10 fell to the same level as that of the control diet despite a 5-fold higher level of serum insulin (Fig 2A, C, Fig 1J, Supp Fig S1H). In muscle tissue, like in eWAT, we found that PTEN increased within 2 weeks (Fig 1O, Supp Fig S2D) and remained significantly increased over 8, 10 (3-fold) and 12 weeks (2.5-fold) in mice on western diet as compared to the control (Fig 2D-E). AKT phosphorylation in mice on western diet progressively lowered by 25% and 50% by weeks 10 and 12 (Fig 2D, F) while the systemic insulin continued to remain 7fold higher (Fig 1J). Also, in support of reduced AKT activity of the muscle, we found that phosphorylated AS160 (readout of glucose uptake and GLUT4 activity in muscle)(*16*) and phosphorylated PRAS40 were also severely suppressed (Supp Fig S2E). Similar observations were made in the liver where at longer times of western diet at 4, 8 and 10 weeks, PTEN was significantly increased by 3.2, 2 and 2.3 folds (Supp Fig S2F-G) while phosphorylated AKT remained the same as control diet at 8 weeks and was lower than control diet by week 10 (Supp Fig S2F, H), in the presence of higher insulin levels in the mice eating western diet.

**Figure 2.**
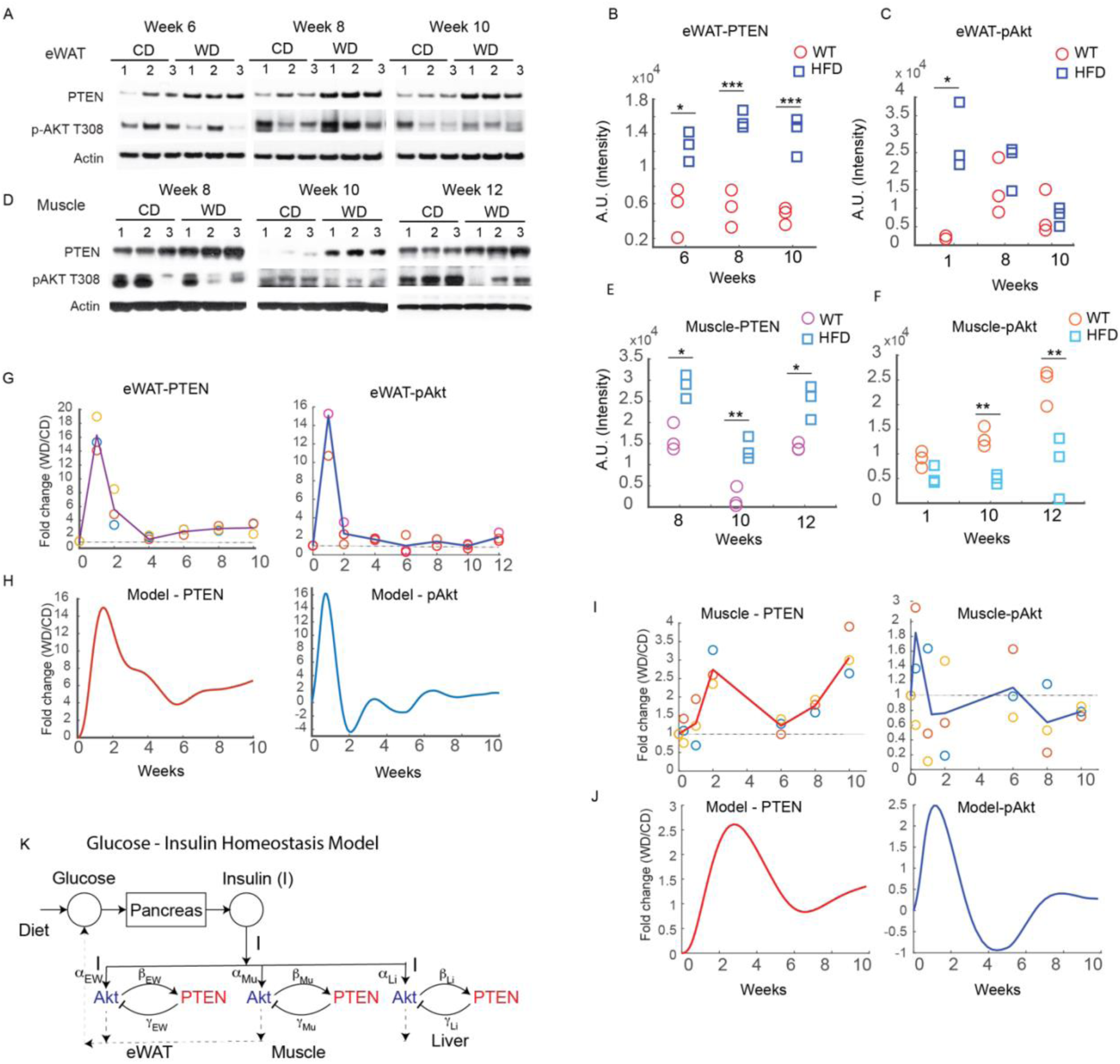
Chronic western diet increases PTEN, decreases AKT activity and is predicted by a computational model. **(A, D)** Mice were either fed with control (CD) or western diet (WD) for 12 weeks, food withdrawn for 3 hours and proteins were extracted from eWAT and muscle and analyzed by western blotting at the indicated times (n=3). **(B-C, E-F)** Quantification of PTEN and pAKT from A, D, p values calculated using student’s t test. *p<0.05, **p<0.01, ***p<0.001. **(G, I)** The PTEN and pAKT signaling intensity of the western blots from A, D and Supp Fig S2C, D were quantified and values in animals on western diet normalized by the values in animals fed control diet and plotted as a function of time. Experimental replicates are indicated in symbols with the mean plotted (n=3). Each symbol represents an animal. **(H, J)** The computational model prediction of the PTEN and AKT values are plotted. **(K)** A computational model of the whole organism glucose-insulin homeostasis was built with systemic glucose(G) driving insulin(I) production which is sensed by the insulin sensitive organs i.e., the eWAT, skeletal muscle and the liver. The insulin drives AKT activation in the tissues which in turn modulates the PTEN levels. PTEN negatively regulates AKT in a negative feedback loop. The effect of the model is to simulate the PTEN and AKT transient behavior as a function of western diet.

Signaling systems that inherently possess negative feedback loops display transient oscillatory behavior seen in the elements of the signaling network(*38–40*). In eWAT we found that both PTEN and pAKT increase together (by 18- and 16-fold respectively) within a week of western diet (Fig 2G, Supp Fig S2C, Fig 1M) in response to a 5 fold increased insulin (Fig 1J). Subsequently both PTEN and pAKT declined and settled at a value where PTEN remained upregulated to about 2-3 fold for 10 weeks and pAKT declined to approximately the same levels as that of the control animals in response to a 5 fold high systemic insulin. There is some transient oscillatory behavior seen in the pAKT and PTEN as a function of time in Fig 2G. Similarly, in skeletal muscle tissue, as seen in Fig 2I, AKT activity peaks at Day 2 and then begins to decrease to a trough by week 2 (Fig 2I, Supp Fig S2D). PTEN is induced starting at 1 week (after the peak of pAKT) and peaks at 2 weeks where it is induced by 2.7 fold over mice on control diet (Fig 2I, Supp Fig S2D). Week 2 is also when severe glucose intolerance sets into the animals on western diet (Fig 1K-L). PTEN decreases from week 2 to week 6 reaching a trough at week 6, while pAKT begins to increase from week 2 and reaches a second peak at week 6. This out of phase behavior continues with PTEN increasing from week 6 and reaching a second peak at 10 weeks (3-fold above control) while pAKT activity decreases to a value of 0.75 of the animals on control diet as shown in Fig 2I. Thus, transient oscillatory behavior in PTEN and pAKT, with pAKT leading PTEN in phase and settling to an attenuated value while PTEN increases to a 2-3 fold higher value in western diet over control diet, is seen in the skeletal muscles.

To quantitatively predict the experimentally observed transient oscillatory behavior that are seen in the different tissues of the animals fed western diet, we built a computational model of the insulin and glucose homeostasis of an animal. As shown in Fig 2K, the glucose in the system post feeding, is sensed and converted into systemic insulin. The insulin is then sensed by the three insulin sensitive peripheral tissues eWAT, skeletal muscles, and liver. The tissues are defined by their PTEN and pAKT values. The insulin drives the pAKT which determines the amount of glucose absorbed by the organ. pAKT drives PTEN while PTEN inhibits pAKT in a negative feedback loop. The same feedback loop scheme is present in every tissue with the difference being that the extent of activation of PTEN, and the extent of negative regulation of pAKT by PTEN, are variables that can differ across the different tissues. The computational model is driven by the systemic glucose and insulin profiles of the animals fed on western diet as the input from Fig 1J-K. This was chosen because while the insulin and glucose profiles of the animals were explainable as consistently increased and many folds higher than control animals due to hyperglycemia and hyperinsulinemia, the PTEN and pAKT profiles are measured to oscillate in different tissue with different kinetic profiles. We varied the parameters of the model, the activation of PTEN by pAKT and the pAKT inhibition by PTEN, in eWAT, muscle and liver respectively. The model parameters were varied until the pAKT, PTEN profiles of the three tissues matched the experimental data using the divided-rectangle optimization and minimization algorithm. A simple model connecting pAKT and PTEN in a negative feedback loop (Fig 2H, J, Supp Fig S2K) for the three tissue was enough to explain the transient oscillations seen in the three tissues with different time constants as is expected with the molecular stoichiometry in the insulin receptor signaling network. Therefore, as hypothesized earlier, initial increase in AKT activity and PTEN levels over chronic exposure to western diet induced hyperinsulinemia leads to a new steady state of heightened negative feedback (PTEN) and reduced AKT activity over time. Transient oscillatory behavior is observed in the kinetics of PTEN and pAKT which is a hallmark of signaling network with feedback loops.

### Inhibiting PTEN activity prevents and reverses insulin resistance

Since PTEN levels increased in insulin sensitive tissues and the increase correlated with the onset of hyperinsulinemia, hyperglycemia and glucose intolerance along with the decline of AKT phosphorylation, we asked whether inhibiting PTEN activity would affect these phenotypes. To that effect we used an inhibitor of PTEN phosphatase activity, VO-OHpic, a vanadyl compound complexed to hydroxypicolinic acid(*41–43*). This complex is a non-competitive inhibitor of PTEN protein and its selectivity towards PTEN over other cysteine-based phosphatases (CBP), including protein tyrosine phosphatases, is based on exploitation of the differences in the catalytic pockets of the phosphatases. The catalytic pocket of PTEN (8Ang) is much larger than that of the other CBPs allowing this compound to bind PTEN with an IC50 of 35nM while it binds other CBPs in the uM range^31^. Its selectivity has been confirmed by its induction of AKT phosphorylation in cells with WT PTEN but not in PTEN null cells^31^. This compound has been tested *in vivo* in multiple pre-clinical models in which it has effectively inhibited PTEN without noticeable toxicity(*41, 44–46*). We confirmed that VO-OHpic induced the PI3K/AKT pathway by treatment of *PIK3CA* mutant BT474 and MCF7 breast cancer cells with the drug (Supp Fig S3A-B). Cancer cell lines with activating mutation in *PIK3CA* and wild type for *PTEN* have higher PTEN expression levels since oncogenic activation of PI3K leads to an increase in PTEN translation(*26*). Upon treatment of these cells with the 1uM VO-OHpic, phosphorylation of AKT, its substrate PRAS40, and mTOR substrates increased as a function of time, consistent with its inhibition of PTEN (Supp Fig S3A-B). In 3T3L-1 adipose cells, VO-OHpic increased the duration of insulin induction of AKT, and S6K phosphorylation (Supp Fig S3C).

Treatment of mice on the Western diet with 10mg/kg of VOOH-pic once daily led to complete prevention of weight gain over 6 weeks (Fig 3A, Supp Fig S3D). No toxicity of the drug was observed. Treatment with VO-OHpic completely prevented hyperinsulinemia, hyperglycemia and insulin resistance as demonstrated by glucose tolerance tests done after 4 and 6 weeks of drug treatment (Fig 3B-D, I-J Supp Fig S3E-F). We confirmed that the mice on western diet with and without drug treatment were consuming approximately the same amount of energy (Supp Fig S3G). We also confirmed that treatment with the inhibitor on control diet did not affect the weight or glucose levels of mice (Supp Fig S3H-I). We asked whether inhibiting PTEN was able to reverse the phenotypes associated with insulin resistance. Mice were treated with VO-OHpic 2 weeks after initiation of the western diet, at which time hyperinsulinemia, hyperglycemia and glucose intolerance were well established (Fig1H-K). VO-OHpic completely reversed weight gain and hyperinsulinemia within 1 week of PTEN inhibition (Fig 3E-F, I Supp Fig S3D). Hyperglycemia and glucose intolerance were reversed after 2 weeks of inhibitor treatment and insulin sensitivity persisted on therapy and the western diet for four weeks (Fig 3G-H, J Supp Fig S3J-K). PTEN inhibition also prevented and reversed the increase of leptin levels within 2 weeks of drug treatment (Fig 3K, Supp Fig S3L). eWAT adipocytes are the primary source of leptin hormone and expand upon consumption of a high fat diet. We found that the adipocyte area per cell increased upon western diet consumption, and this was both prevented and reversed by inhibition of PTEN (Fig 3L-M, Supp Fig S3M).

**Figure 3.**
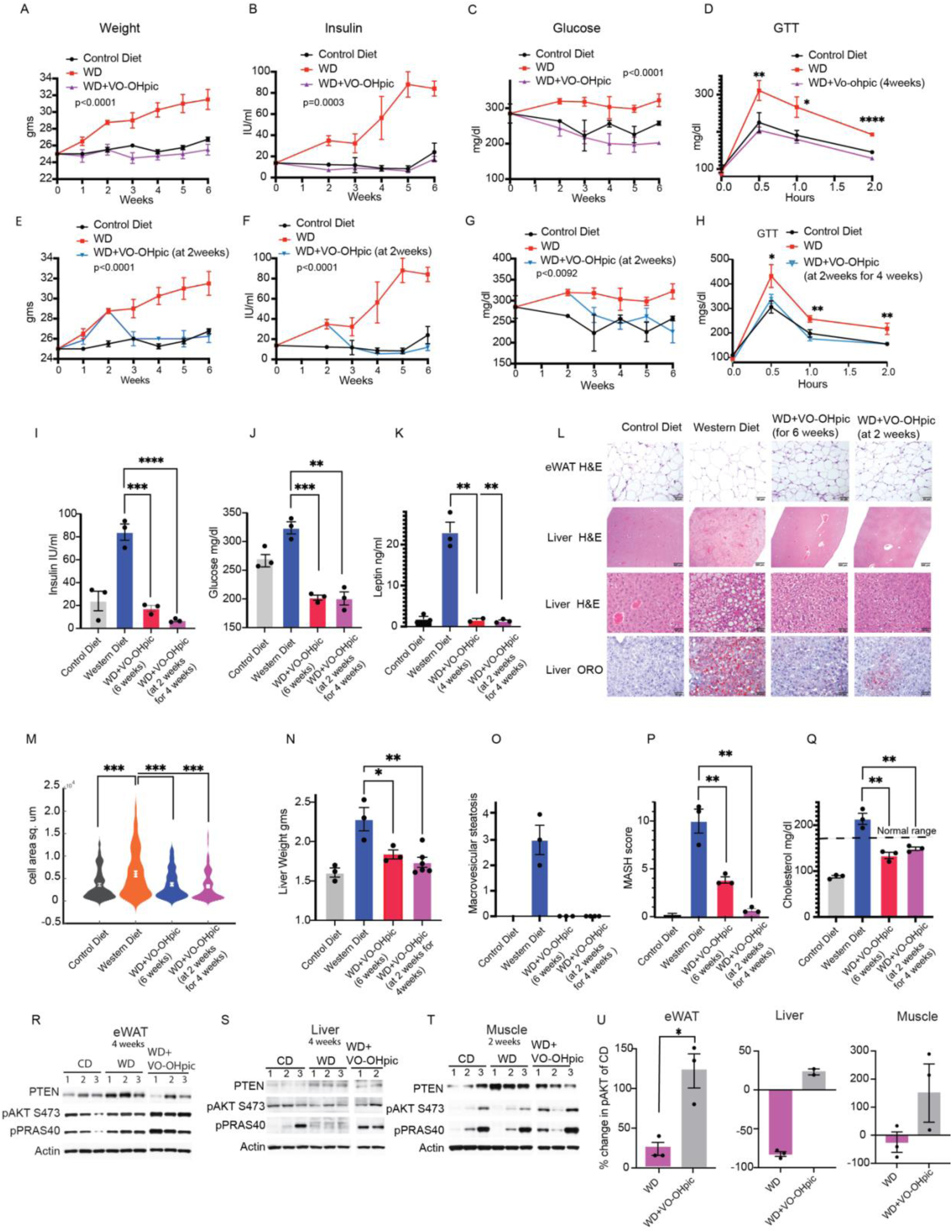
Inhibiting PTEN activity prevents and reverses insulin resistance. **(A-C)** Mice were fed on regular diet or western diet or western diet with treatment of the PTEN inhibitor VO-OHpic (10mg/kg) once daily for 6 weeks and the weight p<0.0001(n=5-10), insulin p=0.0003, glucose p<0.0001 levels measured (n=5). **(D)** Glucose tolerance test (GTT) (n=5) after indicated diet and treatments. **(E-G)** Mice were fed on the indicated diets for 2 weeks and then treated with VO-OHpic (10mg/kg) once daily for 4 weeks and the weight p<0.0001 (n=5-10), insulin p<0.0001 (n=5), glucose levels measured p<0.0092 (n=5). The CD and WD arms of A-C and E-G are the same datasets, since the experiments were done together for consistency. **(H)** glucose tolerance test (GTT) after indicated diets and treatment, n=5. **(I-K)** Insulin (from 3B, F), glucose (from 3C, G) and leptin (K) of mice fed with the indicated diets was measured at the end of the treatment regimens. **(L)** eWAT and livers from mice fed with the indicated diets and treatment stained for H&E and lipid OilRedO (n=3). Two independent experiments done, and representative images are shown. **(M)** Violin plot of morphometric analysis (see Methods) of eWAT H&E from L (n=3). **(N-Q)** The mean and SEM of n was represented of the liver parameters: liver weight, macrovesicular steatosis scores, MASH scores and cholesterol levels (n=3). **(R-T)** eWAT, muscle and liver of mice on control or western diet or western diet treated with VO-OHpic were analyzed by immunoblotting for PTEN and PI3K pathway activity. One of the two independent experiments shown. CD, WD same as Fig 2 since the experimental arms were done together **(U)** Quantification of percent change in pAKT over control from R-T. Mean and SEM of n represented, and p values were calculated by a two-way ANOVA and post Bonferroni tests (A-C, E-G). Mean and SEM plotted, p values calculated by one-way ANOVA test (D, H). Mean and SEM of n represented, p values calculated by student’s t test (I-Q, U).

PTEN inhibition also prevented the increase in liver weight upon western diet and reduced the weight upon inhibitor treatment for 4 weeks after 2 weeks of the diet exposure (Fig 3N). Upon analysis of Oil red O (ORO) staining of liver sections, we found that lipid accumulation was substantially prevented and reversed by inhibiting PTEN at the beginning or after 2 weeks of western diet (Fig 3L). Analysis of H&E sections, that were certified, and blind scored by a pathologist, revealed that VO-OHpic treatment completely prevented and reversed the development of macrovesicular steatosis (within 4 weeks of drug treatment for the latter) (Fig 3O). Microvesicular steatosis was reduced by preventive treatment of VO-OHpic and was reduced more by treating the animals with the drug after 2 weeks of western diet consumption (Supp Fig S3N). Hepatocellular hypertrophy and lobular inflammation both were significantly reduced by preventive treatment while treatment at 2 weeks of the diet completely reversed these conditions (Supp Fig S3O-P). The MASH scores that are an integration of the conditions of macro- and microvesicular steatosis, hepatocellular hypertrophy, and lobular inflammation, averaged around 10 for the mice on western diet, 4 for VO-OHpic treatment at the beginning of the diet and 0.5 for mice that were treated with the inhibitor after 2 weeks on western diet for 4 weeks (Fig 3P). Other markers of liver disease or function (AST, ALT, GGT, albumin) were normal in all groups of mice (data not shown) except cholesterol which was elevated, in the range of hypercholesteremia in mice on the western diet. Treatment with VO-OHpic prevented or lowered the cholesterol levels within the normal range (Fig 3Q). PTEN inhibition for 4 weeks during western diet consumption in the eWAT and liver caused an increase in AKT phosphorylation (80% increase in AKT S473 in eWAT and 105% in liver of mice treated with VO-OHpic over vehicle treated mice on western diet) (Fig 3R-S, U Supp Fig S3Q-R).

PTEN inhibition for as little as 2 weeks in muscle tissue caused an increase of 140% in pAKT S473 in muscle (Fig 3T, U Supp Fig S3S). AKT activity measured by phosphorylated PRAS40 were also increased in all three tissues (Fig 3R-T).

Taken together these data show that inhibition of PTEN phosphatase activity is sufficient to reverse insulin resistance and its metabolic sequelae in mice on a western diet. Insulin induction of PTEN is therefore necessary for maintenance of the phenotype. Moreover, inhibition of PTEN activity in cells in which its expression has been induced by insulin prevents the development of insulin resistance.

### PTEN increase is driven by mTORC1 dependent translation and inhibiting mTORC1 prevents and reverses insulin resistance

PI3K controls mTORC1 regulated 4E-BP1 dependent translation of PTEN protein(*26*). We wanted to investigate if insulin dependent PTEN increase was due to increase in mTOR activity in cells. To that effect we first treated insulin stimulated 3T3L1 cells with mTOR kinase inhibitor AZD8055 that inhibits both mTORC1 and mTORC2 and therefore AKT (pAKT S473 is a mTORC2 target)(*47*). mTOR activity and PTEN expression was increased during insulin stimulation (Fig 4A). In cells co-treated with insulin and AZD8055, mTORC1 and 2 activity was inhibited as measured by the de-phosphorylation of 4E-BP and S6K and AKT S473 while the increase in PTEN protein was completely rescued and it remained reduced over time (Fig 4A). We asked whether selective inhibition of TORC1 kinase prevented the increase in PTEN protein by insulin stimulation. RMC-6272 is a selective inhibitor of TORC1 kinase (IC50 for mTORC1 inhibition is 0.44 nM for p4E-BP, IC50 for TORC2 inhibition of pS473 AKT is 12nM)(*48*). At TORC1 selective doses this drug inhibits 4E-BP1 phosphorylation but not the TORC2 dependent AKT phosphorylation. Treatment with RMC-6272 prevented the induction of PTEN after insulin stimulation in 3T3L1 adipocytes (Fig 4B) and reversed PTEN increase when cells were treated with the drug after 4 hours of insulin stimulation when mTORC1 activity and PTEN levels were allowed to increase first (Supp Fig S4A-B). In insulin treated cells PTEN was increased by approximately 1.8 fold after 48 hours, and when treated with RMC-6272 after 4 hours of insulin treatment, PTEN levels were reduced to 0.8 fold of unstimulated cells. Reduction in PTEN level was accompanied by increased duration of AKT phosphorylation in RMC-6272 inhibitor and insulin treated cells (Supp Fig S4A).

**Figure 4.**
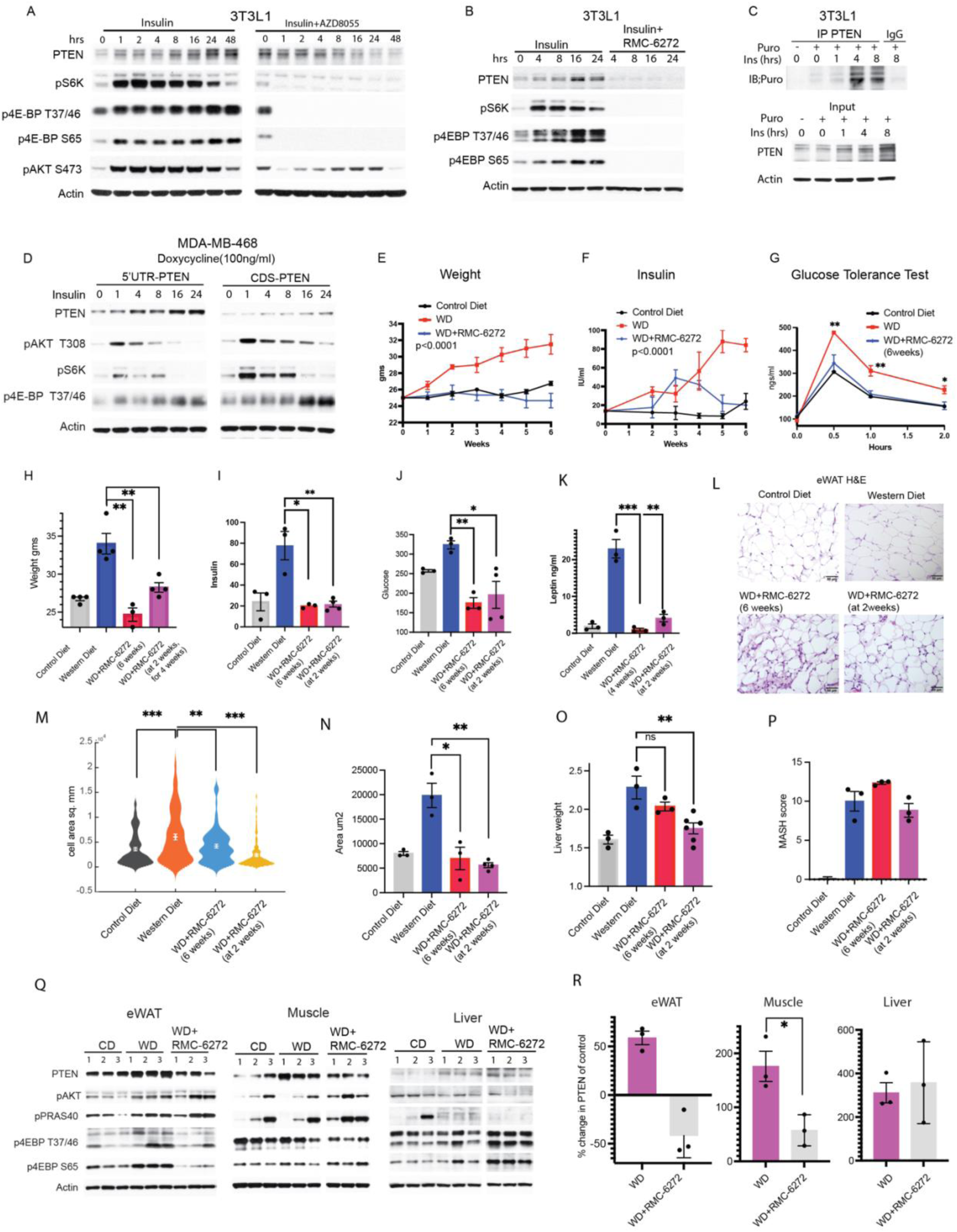
PTEN increase is driven by mTORC1 dependent translation and inhibiting mTORC1 prevents and reverses insulin resistance. **(A)** 3T3L1 cells were treated with 1ug/ml of insulin alone or a combination of insulin and 500nM AZD8055 for the indicated times and analyzed by immunoblotting for PTEN and mTORC pathway activity. **(B)** 3T3L11 cells were treated 1ug/ml insulin alone or in combination with RMC-6272 (0.5nM) for the indicated times and immunoblotted for PTEN and mTORC1 pathway activity. **(C)** 3T3L1 cells were treated with insulin for the indicated times and pulsed with puromycin for 30 min and PTEN was immunoprecipitated and immunoblotting done for puromycin (See Methods). **(D)** MDA-MB-468 cells expressing a doxycycline inducible PTEN with or without its 5’UTR was treated with 100ng/ml of doxycycline for 24 hours and then stimulated with insulin for the indicated times and PTEN protein analyzed by immunoblotting. **(E-F)** Mice were fed on regular diet or western diet or western diet treated with the mTORC1 inhibitor RMC-6272 (3mg/kg) once a week for 6 weeks and the weight p<0.0001 (n=5-10), insulin p<0.0001 (n=5) measured. **(G)** GTT measured at the indicated time, diet and treatment conditions (n=5). **(H-K)** Weight (from 4E, Supp S4H), Insulin (from F, Supp S4I), glucose (from Supp S4D, J) and leptin of mice fed with the indicated diets were measured at the end of the treatment regimens. **(L)** eWAT from mice treated with indicated diet and treatment conditions, stained for H&E, representative images shown (n=3). Two independent experiments were done, and representative images are shown. **(M-N)** Violin plot of morphometric analysis of eWAT H&E from L (n=3). N. Mean and SEM of n of morphometric analysis from M was plotted. **(O-P)** Liver weight and MASH scores were measured at the indicated times and treatment conditions (n=3), also see from Supp S4M. **(Q)** eWAT, liver and muscle of mice that were fed with indicated diets and treatment conditions were analyzed for PTEN and PI3K pathway activity by immunoblotting, n=3. CD, WD same as Supp Fig 2 since the indicated experimental arms were run on the same gels together **(R)** Quantification of percent change in PTEN over control diet from Q. Mean and SEM of n represented, and p values were calculated by a two-way ANOVA and post Bonferroni tests (E-F). Mean and SEM plotted, p values calculated by one-way ANOVA test (G). Mean and SEM of n represented, p values calculated by student’s t test (H-K, M-P, R).

Next, we wanted to investigate if mTOR dependent increase in PTEN during insulin stimulation was due to increase in its protein translation. We found that when we treated the cells with global translational inhibitors like cycloheximide and eIF4A inhibitor, Silvestrol(*47*), insulin dependent PTEN protein increase was rescued and PTEN expression was reduced (Supp Fig S4C). We directly confirmed that PTEN translation increased during insulin stimulation by labelling with Puromycin, an aminonucleoside antibiotic that incorporates in the C terminus of elongating amino acid chains(*49*). We stimulated 3T3L1 cells with insulin for 1, 4 and 8 hours with insulin and pulsed with puromycin for 30 minutes and pulled down PTEN from protein lysates and blotted for puromycin. We found that the puromycin signal was upregulated at 4 and 8 hours suggesting an increase in de novo translation of PTEN during insulin stimulation (Fig 4C). mTOR phosphorylates 4E-BP and causes it to dissociate from the translation initiation factor eIF4E, thus relieving its inhibition of formation of the eIF4F initiation complex. This allows the translation initiation complex to scan the 5’UTR of capped mRNAs permitting translation initiation(*30, 31*). In MDA-MB-468 PTEN null cells, expression of PTEN coding sequence with the 5’UTR (that allows for its translational control) showed increase in PTEN protein during insulin stimulation whereas expression of PTEN without its 5’UTR (unregulatable by translation(*26*)) prevented the induction of PTEN expression during insulin stimulation. The duration of insulin stimulated AKT phosphorylation and downstream pathway activation (pS6K and p4EBP) was increased in cells that expressed 5’UTR less PTEN where PTEN levels did not increase as compared to those in which PTEN increased substantially (Fig 4D). Taken together these experiments suggest that insulin stimulation increases mTORC1 dependent PTEN translation and that this mechanism is an important regulator of duration of AKT signal during insulin stimulation.

We wished to confirm that the diet dependent increase in PTEN protein in mice is sensitive to mTORC1 inhibition. We tested whether RMC-6272 inhibited the increase in PTEN expression that occurs in mice on the western diet and, in doing so, prevents its induction of obesity, insulin resistance and MASH. RMC-6272 (3m/kg) was administered to mice once per week when the western diet was initiated. Treatment of RMC-6272 at the beginning of the diet completely prevented weight gain, reduced insulin levels after 4 weeks and glucose levels after 2 weeks to normal levels and prevented the development of insulin resistance (Fig 4E-G, H-J, Supp Fig S4D). We confirmed that the mice in different groups approximately had the same energy intake (Supp Fig S4E, Methods) and the inhibitor alone did not induce major changes in weight or glucose levels in animals (Supp Fig S4F-G).We found that weight gain, hyperinsulinemia, hyperglycemia and the development of insulin resistance were completely reversed when the mice were treated with the drug two weeks after the diet was given to them (Supp Fig S4H-K, Fig 4H-J). mTORC1 inhibition also prevented and reversed increases in leptin levels (Fig 4K, Supp Fig S4L) and adipocyte cell area in eWAT tissues (Fig 4L-N).

By contrast, although PTEN inhibition prevented and reversed the MASH phenotypes in mice fed on the western diet, mTORC1 inhibition did not. Liver weight, lipidosis, steatosis and overall MASH scores were not decreased by RMC-6272 (Fig 4O-P). This result was confirmed by H&E and ORO staining of liver sections that showed no reduction in the lipid accumulation had occurred on the western diet (Supp Fig S4M). Neither was the diet-induced hypercholesterolemia reduced by mTORC1 inhibition (Supp Fig S4N) whereas serum liver enzymes such as serum AST, ALT, GGT as well as albumin levels were normal in all groups of mice (data not shown). To confirm that the failure to inhibit the MASH phenotype was not specific to RMC-6272, we used a different mTORC1 inhibitor, Rapamycin(*50*). Rapamycin inhibited weight gain in mice on western diet and quite effectively reduced hyperinsulinemia and hyperglycemia (Supp Fig S4O-Q). However, H&E sections of the liver treated with Rapamycin revealed the presence of liver lipidosis (Supp Fig S4R).

Lastly, we analyzed pAKT/mTORC1 activity and PTEN protein levels in the eWAT, liver and muscle tissues upon treatment with RMC-6272 or Rapamycin. Treatment of RMC-6272 or Rapamycin along with consumption of western diet for 4 weeks in the eWAT or 2 weeks in the muscle, led to a decrease in p4E-BP1 T47/46 and S65 sites and there was a decrease in PTEN expression accompanied by increase in pAKT (Fig 4Q-R, Supp Fig S4S). In contrast, in the liver, neither RMC-6272 nor Rapamycin treatment for 4 weeks inhibited mTORC1 activity when the mice were fed with western diet, and PTEN levels were almost unchanged (Fig 4Q-R, Supp Fig S4S). Consistent with persistent PTEN overexpression, pAKT T308 phosphorylation and that of the AKT substrate remained low (Fig 4Q).

In summary, increase in PTEN during insulin stimulation is regulated by mTORC1 mediated cap-dependent translation. Inhibiting PTEN activity in the insulin sensitive tissues inhibited the obesity, insulin resistance and MASH phenotypes whereas inhibiting mTORC1, suppressed PTEN and its metabolic effects in eWAT and muscle but not liver. The difference seems to be the effectiveness of the mTORC1inhibitors to reduce PTEN in the liver.

## Discussion

Insulin resistance and subsequent type 2 diabetes are complex phenomena for which there is currently no unitary explanation(*51*). No single primary event that affects the insulin signaling pathway and explains multiple features of the diseases has been found, although in single cases and in families with inherited disorders of glucose homeostasis it has been shown to be due to single gain or loss of function mutations that affect signaling (e.g. insulin receptor, IRS proteins, TSC2, AKT2)(*52–54*). Various phenomena that are part of the syndrome (increased lipolysis in fat cells, increased gluconeogenesis and glycogenesis in liver) have been known to cause other aspects of the phenotype(*16*). Changes in neural and endocrine regulation of appetite have been shown to be associated with the syndrome but beg the question of how increased food intake initiates the problem(*2, 4*).

We show here a relatively simple mechanism that explains insulin resistance with attendant effects on white adipose, liver, and muscle cells that explain many of the features of type two diabetes. It is based on a recent finding that the translation of PTEN, a potent downstream negative regulator of insulin signaling, is mTOR dependent(*26*). Activation of the insulin/PI3K/AKT/mTOR pathway is therefore buffered by induction of PTEN expression. Similarly, nutrient deprivation or reduction in insulin signaling causes a fall in PTEN expression, allowing of some level of AKT signaling. Hence, PTEN is a powerful feedback regulator of insulin signaling that is predicted to play an important role in metabolic homeostasis.

This model generated the hypothesis that hyperactivation of insulin could lead to an overshoot of PTEN expression and thus insulin resistance. We show here that this is the case in mice on a high fat and carbohydrate diet. After initiation of the diet, insulin levels increase and PTEN expression increases in white adipose tissue, muscle and liver. This is followed by weight gain, hyperglycemia, insulin resistance, hepatic lipidosis, and leptin resistance. In support of our hypothesis, each of these is prevented or reversed by administration of a selective inhibitor of the PTEN lipid phosphatase. Also, tissue specific PTEN knockout models generated by others in the field have shown improved insulin sensitivity and its associated metabolic phenotypes(*29*). Caloric restriction is also expected to inhibit mTOR and thereby inhibit PTEN. This may explain in part why it is effective in treating insulin resistance and T2D(*55*). It has also been reported that 4E-BP1 and 4E-BP2 double knockout mice have increased sensitivity to obesity and insulin resistance whereas overexpressing 4E-BP1 makes mice resistant to the phenotypes(*56, 57*). This could be explained by increased translation of PTEN protein in 4E-BP knockout mice and its decrease in the overexpression models.

PTEN inactivates AKT signaling by dephosphorylating PIP3(*25*). PTEN is both a lipid and protein phosphatase and was previously shown to dephosphorylate the tyrosine 612 residue on IRS1 leading to IRS1 and then AKT activations(*58*). Thus, induction of PTEN by the insulin pathway may cause feedback inhibition of the pathway by two mechanisms. This model may explain the loss of IRS1 activity in mice with insulin resistance and in women diagnosed with gestational diabetes that has been reported before(*59–61*).

The complex relationship between AKT and PTEN is attributed to them being connected in a negative feedback loop. In response to hyperinsulinemia both PTEN and AKT transiently oscillate in the insulin sensitive peripheral tissues before settling into a steady state of prolonged insulin resistance. While theoretically the experimental verification of the transient oscillations would require infinitely many time sampling points with multiple animals per point, the use of a computational model to predict the dynamics ameliorates the need for increased number of animals in the experiments.

This paper reports a potential mechanism for development of insulin resistance and diabetes, but it may also have therapeutic implications. In animals treated with a selective PTEN inhibitor, insulin resistance and some of its key biological consequences are prevented and reversed, suggesting the potential use of such a drug in patients. We do not know the long-term consequences of taking a PTEN inhibitor. PTEN is a tumor suppressor gene and loss of a single copy may be haploinsufficient. It may be possible to determine doses that inhibit elevated activity of the protein but do not cause inhibition below physiologic levels. This has not yet been tried. Another possibility is the use of mTOR inhibitors which reduce PTEN translation and expression. However, although we found that mTORC1 selective inhibitors reverse obesity and insulin resistance in mice on the Western diet they do not reverse MASH. This is associated with desensitization of liver mTORC1 to these drugs. Moreover, mTORC1 inhibition with Rapalogs have been observed to induce paradoxical insulin resistance(*62*).

## Acknowledgements

We thank the Anti-tumor assessment core, the Laboratory of Comparative Pathology core for assistance with animal necropsies and histopathology experiments.

## Author Contributions

Conceptualization: RM, NR

Methodology: RM, PP, MS, HS, MT, CT, OH, VM, JD, SM, JC, SEC, ES, KGV

Investigation: RM, SEC, KGV, NR

Funding acquisition: NR

Project administration: RM, MT, NR

Supervision: RM, NR

Writing –RM, KGV, NR

## Data and materials availability

All raw data, code, and materials used will be available to any researcher upon asking.

## Supplementary Materials

### Materials and Methods

#### Animal studies-weight measurements and blood parameters

For the western diet induced obesity and insulin resistance phenotypes, C57BL/6J mice at 8 weeks of age were placed on either a standard laboratory rodent chow or western diet (D12079B, Research Diets) and allowed to eat *ad libitum* and their weights measured as indicated. Food withdrawn for 3 hours, and blood serum was collected and measured for glucose, cholesterol and liver function tests using chemical analyzers and insulin and leptin using ELISA at the indicated times.

#### Glucose tolerance test

Mice were fasted for overnight, and glucose (2g/kg) was intraperitoneally injected into the mice and blood collected from the tail vein and glucose concentrations were determined at the 30minutes, 1hr and 2hrs.

#### Tissue collection and western blotting

Mice were allowed to eat *ad libitum* and food withdrawn for 3 hours and epididymal white adipose tissue, liver, leg and arm muscles were collected from mice at the end of each time point (as indicated) and flash frozen. They were homogenized in SDS lysis buffer (50mM Tris-HCL pH 7.4, 10% Glycerol, 2% SDS) and boiled at 95 C for five minutes. Lysates were then briefly sonicated, boiled again for 5 minutes, before clearing by centrifugation at 14,000rpm for 10 minutes at room temperature. The supernatant was collected, and protein concentration was determined using the BCA kit (Pierce) per manufacturer’s instructions. Protein samples were diluted in SDS sample buffer (final concentration: 62.5mM TrisHCL pH 6.8, 2% SDS, 10% Glycerol, 15.5mg/mL DTT, 0.02mg/mL Bromophenol blue). 25-50 mg of protein was loaded onto each lane of a 4%–12% BisTris mini gel or midi gel (Invitrogen) for immunoblotting. Transfer was onto nitrocellulose membranes (0.2 mm, GE Health Care) before blocking for 1h at room temperature and incubating with primary antibodies of the indicated protein targets overnight at 4 C. Membranes were incubated with secondary rabbit antibody (Sigma) or secondary mouse antibody (GE Health Care) for 1h at room temperature. Blots were developed in Perkin-Elmer’s Western Lightning ECL or Millipore’s Immobilon HRP reagents per manufacturer’s instructions.

### Cell lines and Antibodies and drugs

3T3L1, C2C12, HEPG2, BT474, MCF7, MDA-MB-468, CHO cells were acquired from ATCC and cultured in DMEM-F12 and in DMEM (refer ATCC). All cell lines were supplemented with 10% Fetal Bovine serum (FBS) and 1% penicillin and streptomycin and 4mM Glutamine.

Antibodies used are PTEN (CST #9559), pAKT T308 (CST #2965), pAKT S473 (CST #4060), pPRAS40 (CST #2997), p4E-BP1 T37/46 (CST#2855), p4E-BPs S65 (CST #9451), IRS1 (CST# 2382), IR-ß (CST #3025), ß-Actin (CST #4970)

### Histological analysis

Representative sections of the liver, pancreas, brain, epididymal adipose tissue, retroperitoneal white adipose tissue, skeletal muscle from forelimbs, and skeletal muscle from the hindlimbs were fixed in 10% neutral-buffered formalin, processed in alcohol and xylene, embedded in paraffin, sectioned (5-μm-thick) and stained with hematoxylin and eosin. Oil red O staining was performed on formalin fixed, OCT-embedded frozen sections (5-μm-thick) of liver. For histopathological analysis, hematoxylin–eosin-stained or ORO-stained tissue specimens were evaluated by a board-certified veterinary pathologist (S.E.C.). Liver sections were evaluated and scored, using a semiquantitative histopathology scoring system, with slight modifications, for mouse model of metabolic dysfunction associated fatty liver disease(*63*). Briefly, macrovesicular steatosis, microvesicular steatosis and hepatocellular hypertrophy were separately scored, and the extent and severity of the lesions were graded, into the following categories: 0 (<5%), 1 (5-10%), 2 (10–25%), 3 (25-75%) and 4 (>75%). Inflammation was evaluated by counting the number of inflammatory foci per five 100x fields using the following categories: normal (<0.5 foci), minimal (0.5-1.0 foci), mild (1.0-2.0 foci), moderate (2.0-5.0 foci), severe (>5.0 foci). An Olympus BX45 light microscope was used to capture images with a DP26 camera using cellSens. Dimension software (v1.16).

### Immunohistochemistry

Immunolabeling of PTEN in liver and epididymal white adipose sections was performed at the MSK Biobank and Pathology Core facility. Formalin-fixed, paraffin-embedded sections were stained using an automated staining platform. Briefly, following deparaffinization and heat-induced epitope retrieval, the primary antibody against PTEN (1:200, Cat. No 9559, clone 138G6, Cell Signaling Technologies) (*64*).

### Computational Model Methods

As shown in Fig 2I, the computational model is a model for the whole body glucose and insulin homeostasis in animals fed high fat and high carbohydrate diet (western diet). The computational model features systemic Glucose (G), insulin (I), PTEN and AKT for the three insulin sensitive peripheral tissues of adipose, skeletal muscle and liver respectively.

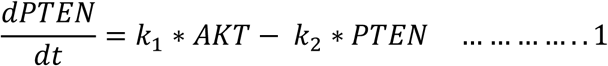

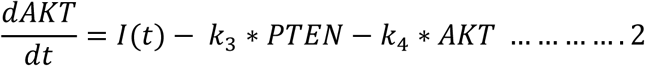

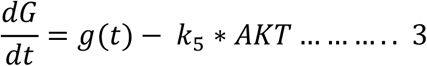

Where *k*_1_ is the feedback factor and defines the extent of activation of PTEN by AKT in the feedback loop. *k*_2_is a negative factor that degrades PTEN. This represents endogenous PTEN degradation by cell intrinsic mechanisms. *k*_3_is the factor for the negative regulation of PTEN on AKT. *k*_4_ is the negative effect of the phosphatases and other deactivators of AKT and term is proportional to AKT. *k*_5_determines the rate at which AKT activation results in the absorption of glucose into the cell and a reduction in the systemic glucose. The equations were implemented in MATLAB and the parameters were estimated in a divided rectangle parameter search method.

### Puromycin pulse assay for *de novo* translation

3T3L1 cells were treated with 1ug/ml of insulin and pulsed with 1uM of puromycin for 30 minutes, collected, lysed, protein extraction and immunoprecipitation of PTEN done using PTEN antibody conjugated to Sepharose beads (CST #4326) or IgG control and immunoblotting done for puromycin (Kerafast #EQ0001) using the above western blotting technique

### Morphometric analysis of eWAT

Cell size distribution in hematoxylin-eosin (H&E)-stained sections of epididymal white adipose tissue was analyzed from triplicates of 40X images per group and cell size was quantified using Adiposoft Software (Image J)(*65*)

### Treatment with PTEN inhibitor VO-OHpic

VO-OHpic was suspended in 2% DMSO, 40% PEG 300, 5% Tween-80, ddH2O and administered intraperitonially at a dose of 10mg/kg, every day, once a day. This was done either at the same time as the start of the western diet in mice for 6 weeks or after 2 weeks of western diet feeding for 4 weeks

### Treatment with mTORC1 inhibitor RMC-6272 or Rapamycin

RMC-6272 was suspended in 1:1 (v/w) Transcutol/Solutol HS 15 and administered intraperitonially at a dose of 3mg/kg, once a week. This was done either at the same time as the start of the western diet in mice for 6 weeks or after 2 weeks of western diet feeding for 4 weeks. Rapamycin was dissolved in 100% DMSO and administered intraperitonially at a dose of 10mg/kg, three times a week along with western diet.

### Other drugs used

Cycloheximide, Silvestrol, AZD8055 were acquired from Selleckchem.

---

**Supplemental Fig S1.**
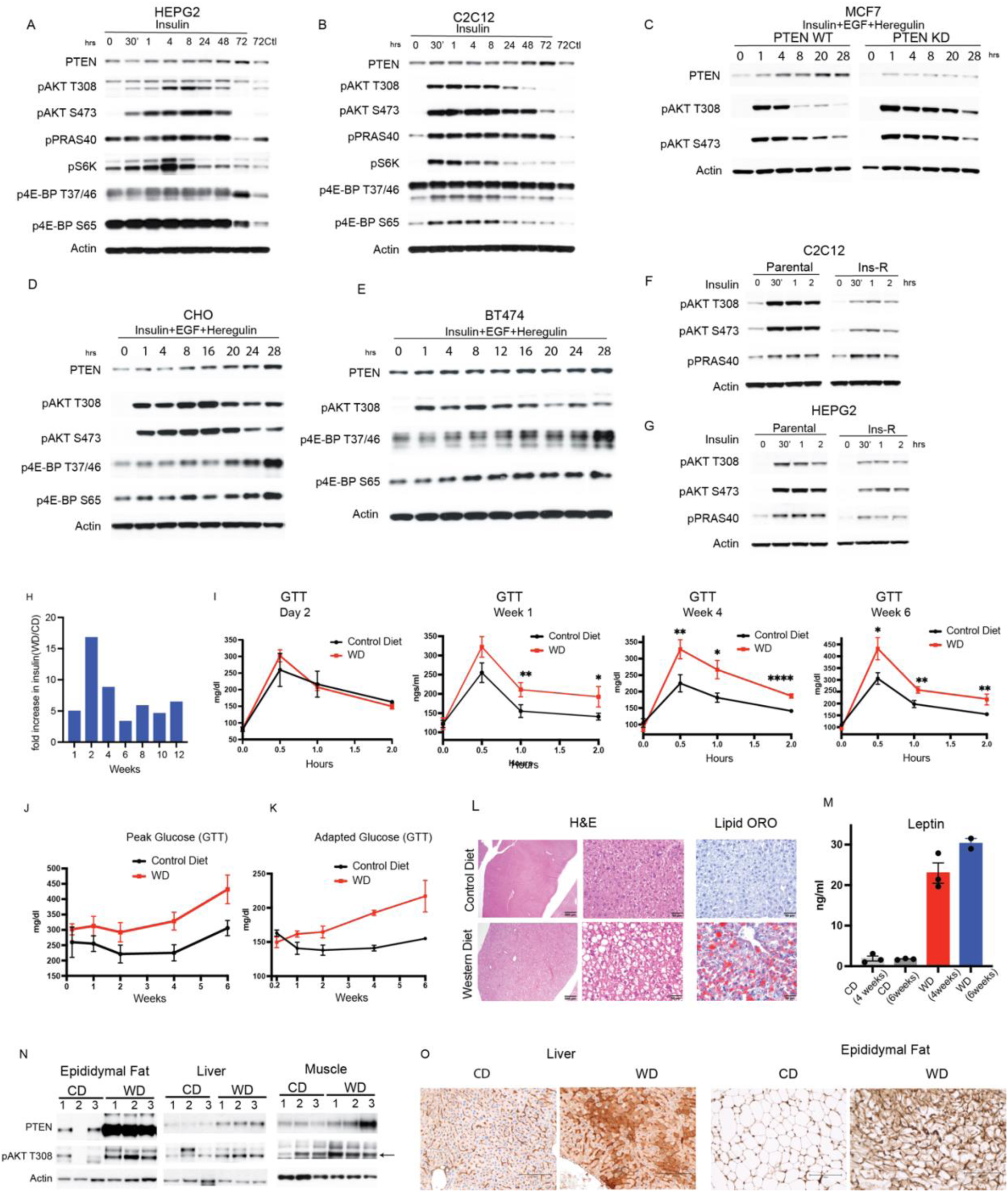
Western Diet in mice and insulin stimulation in cells increase PTEN expression. **(A-B)** HEPG2 and C2C12 cells were treated with 1ug/ml insulin and analyzed for PTEN and PI3K pathway activity by immunoblotting. **(C-E)** CHO, BT474 and MCF7 (PTEN WT and knockdown) cells were treated with a cocktail of Insulin, EGF and Heregulin at 100ng/ml each for the indicated times and analyzed for PTEN and PI3K pathway activity by immunoblotting. **(F-G)** Immunoblotting of Par or Ins-R cells, washed out after 7 days of insulin treatment, and re-treated with 100ng/ml of insulin. **(H)** Fold increase in insulin of C57BL/6J mice fed with western diet over mice fed with control diet for 12 weeks, from Fig 1J (n=5). **(I)** Glucose Tolerance Test (GTT) was performed 2 days, 1, 2, 4 and 6 weeks after start of the diet (n=5) **(J)** Glucose levels at 30 min (peak glucose) of GTT from I and Fig 1L was plotted over time for mice on control or western diet. **(K)** Glucose levels at 2 hours (adapted glucose) of the GTT from I and Fig 1L was plotted over time for mice on control or western diet. **(L)** Mice were either fed with control or western diet for 4 weeks and livers stained for H&E and Oil Red O (ORO) (n=3). Representative images of one of two independent experiments shown. **(M)** Leptin levels were measured in mice after 4 and 6 weeks of control and western diet (n=3) **(N)** Mice were either fed with control (CD) or western diet (WD) and proteins were extracted from eWAT, liver and muscle and analyzed by immunoblotting (n=3, also refer Fig 1M-O). **(O)** Liver and eWAT tissues from mice fed on CD or WD were immunostained for PTEN and representative images shown (n=3), one of two independent experiments represented, also see Fig 1Q). Graphs were plotted as SEM of n, and p values were calculated by Student’s unpaired t test. *p<0.05, ** p<0.01, **** p<0.0001.

**Supplemental Figure S2.**
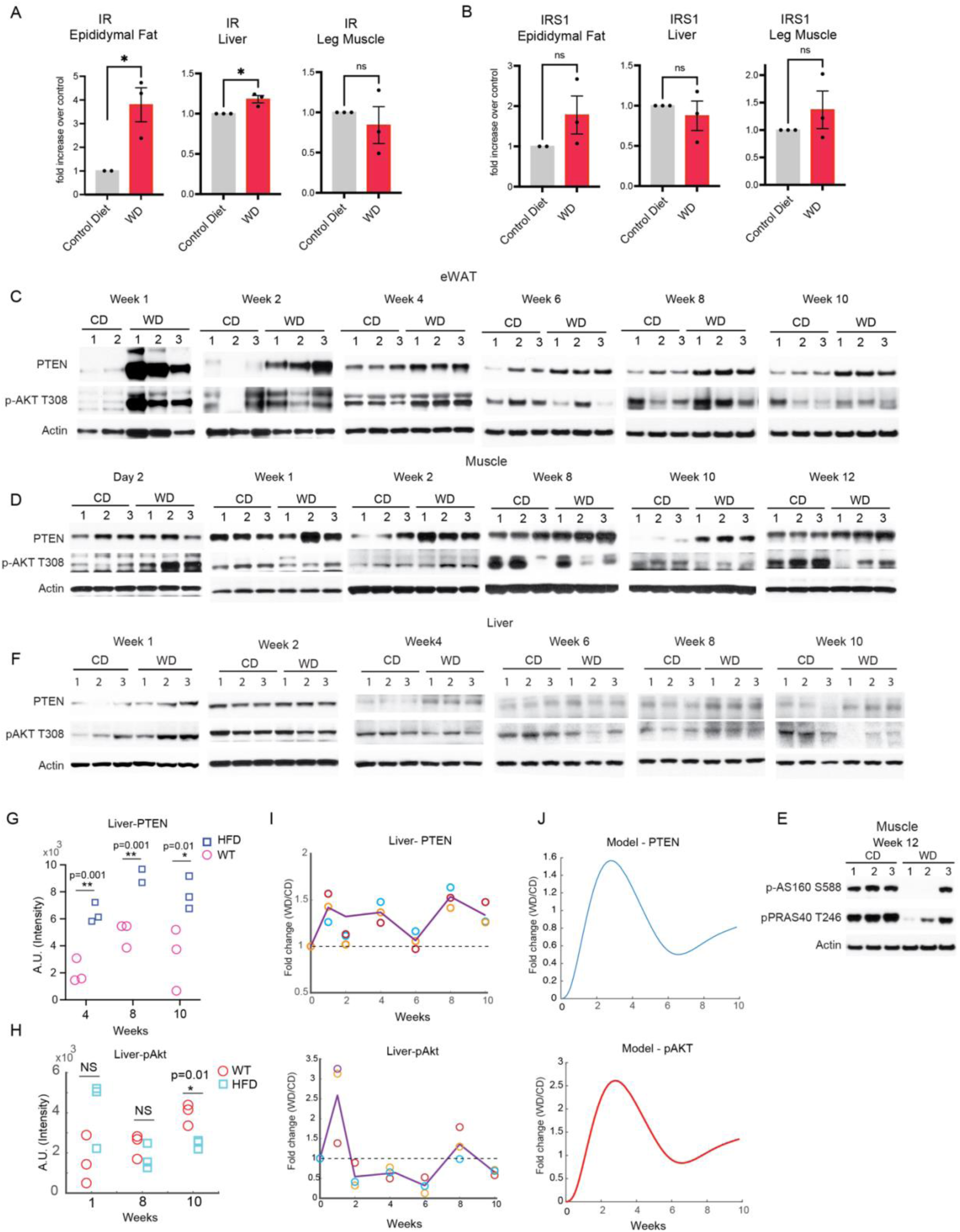
Chronic western diet increases PTEN, decreases AKT activity and is predicted by a computational model. **(A-B)** IR and IRS1 from Fig 1M-O was quantified and normalized with Actin and the mean and S.E.M represented (n=3). p value was calculated using student’s t test. **(C-F)** Mice were either fed with control (CD) or western diet (WD) for 12 weeks, food withdrawn for 3 hours and proteins were extracted from eWAT, muscle and liver and analyzed by western blotting at the indicated times (n=3). **(G-H)** Quantification of PTEN and pAKT from liver (F), p values calculated using student’s t test (n=3) *p<0.05, **p<0.01, NS=non-significant. **(I)** The PTEN and pAKT signaling intensity of the western blots from the liver samples (F) and were quantified and values in animals on western diet normalized by the values in animals fed control diet and plotted as a function of time. Different experimental replicates are indicated in symbols with the mean plotted (n=3). Each symbol represents an animal. **(J)** The computational model prediction of the PTEN and pAKT values are plotted.

**Supplemental Figure S3.**
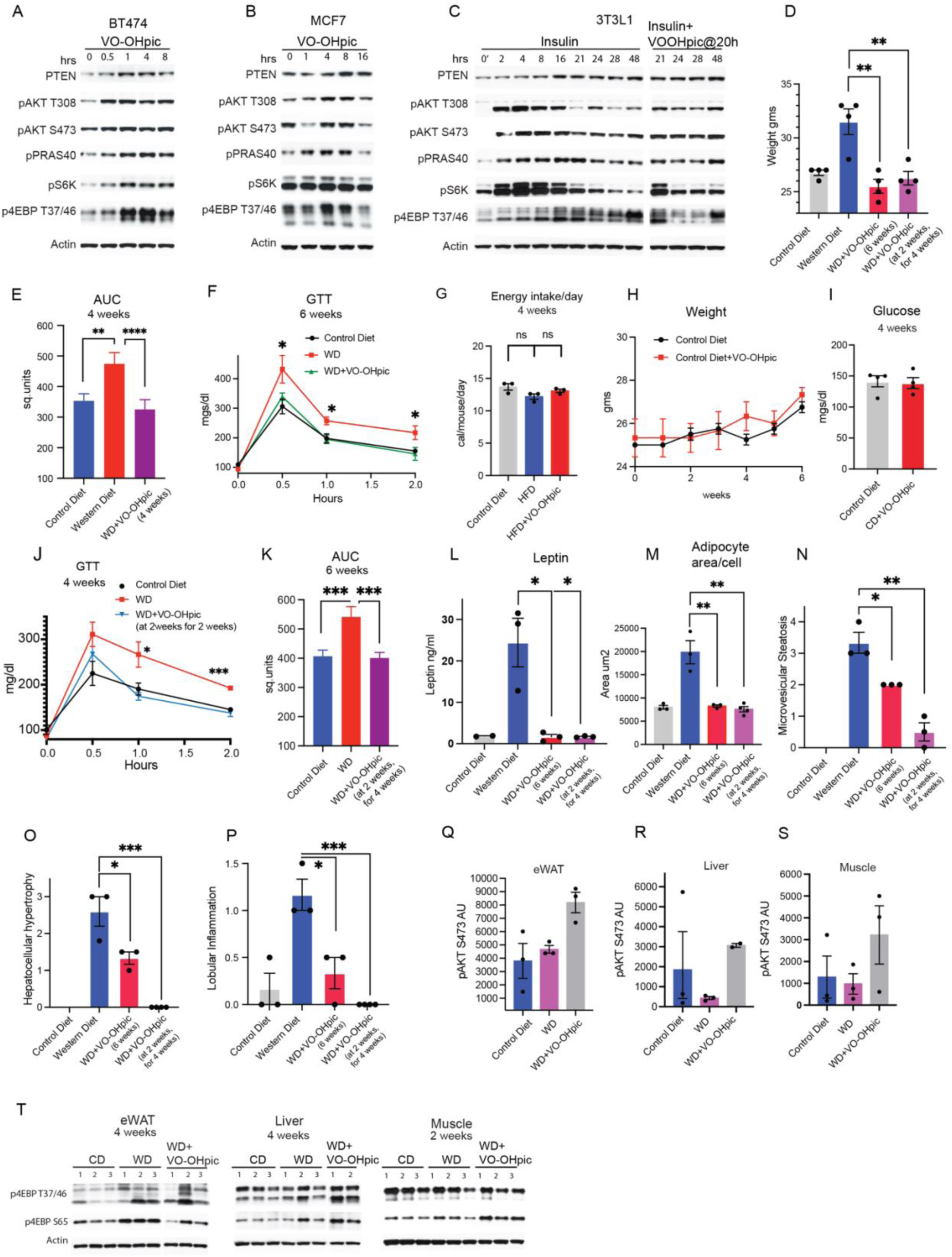
Inhibiting PTEN activity prevents and reverses insulin resistance. **(A-B)** BT474 and MCF7 cells were treated with 1uM VO-OHpic for the indicated times, protein extracted and immunoblotted for PTEN and PI3K pathway activity. **(C)** 3T3L1 cells were treated with1ug/ml of insulin alone or with insulin and VO-OHpic (1uM) added at 20 hours of insulin treatment and immunoblotted for PTEN and PI3K pathway activity. **(D)** Weight of mice fed with the indicated diets at the end of the treatment regimens was measured (from Fig 3A, E), (n=5-10). **(E)** Area under the curve (AUC) was calculated of the GTT assay (Fig 3D) for the indicated times and conditions of diet and VO-OHpic treatment. **(F)** GTT measured for the indicated times and treatment conditions, (n=5). **(G-I)** Mean and SEM of energy intake, weight and glucose measured of mice for the indicated times and conditions (n=5). **(J)** GTT measured for the indicated times and treatment conditions, (n=5). **(K)** Area under the curve (AUC) was calculated of the GTT assay (Fig 3H) for the indicated times and conditions of diet and VO-OHpic treatment. **(L)** Leptin was measured for the indicated times and treatment conditions (n=3). **(M)** Area of eWAT adipocyte cells from Fig 3L-M was measured for the indicated dietary and treatment conditions (n=3). Two independent experiments done and representative images shown. **(N-P)** The mean and SEM of liver parameters microvesicular steatosis, hepatocellular hypertrophy, and lobular inflammation scores measured and mean and SEM of n represented (n=3). **(Q-S)** Quantification of pAKT S473 normalized to actin from Fig 3R-T, mean and SEM of n plotted. **(T)** eWAT, muscle and liver of mice on control or western diet or western diet treated with VO-OHpic were analyzed by immunoblotting for mTORC1 activity. es calculated by student’s t test (D-E, K-P). Mean and SEM of n plotted, p values calculated by one-way ANOVA test (F, J). *p<0.05, **p<0.01, ***p<0.001, ****p<0.0001.

**Supplemental Figure S4.**
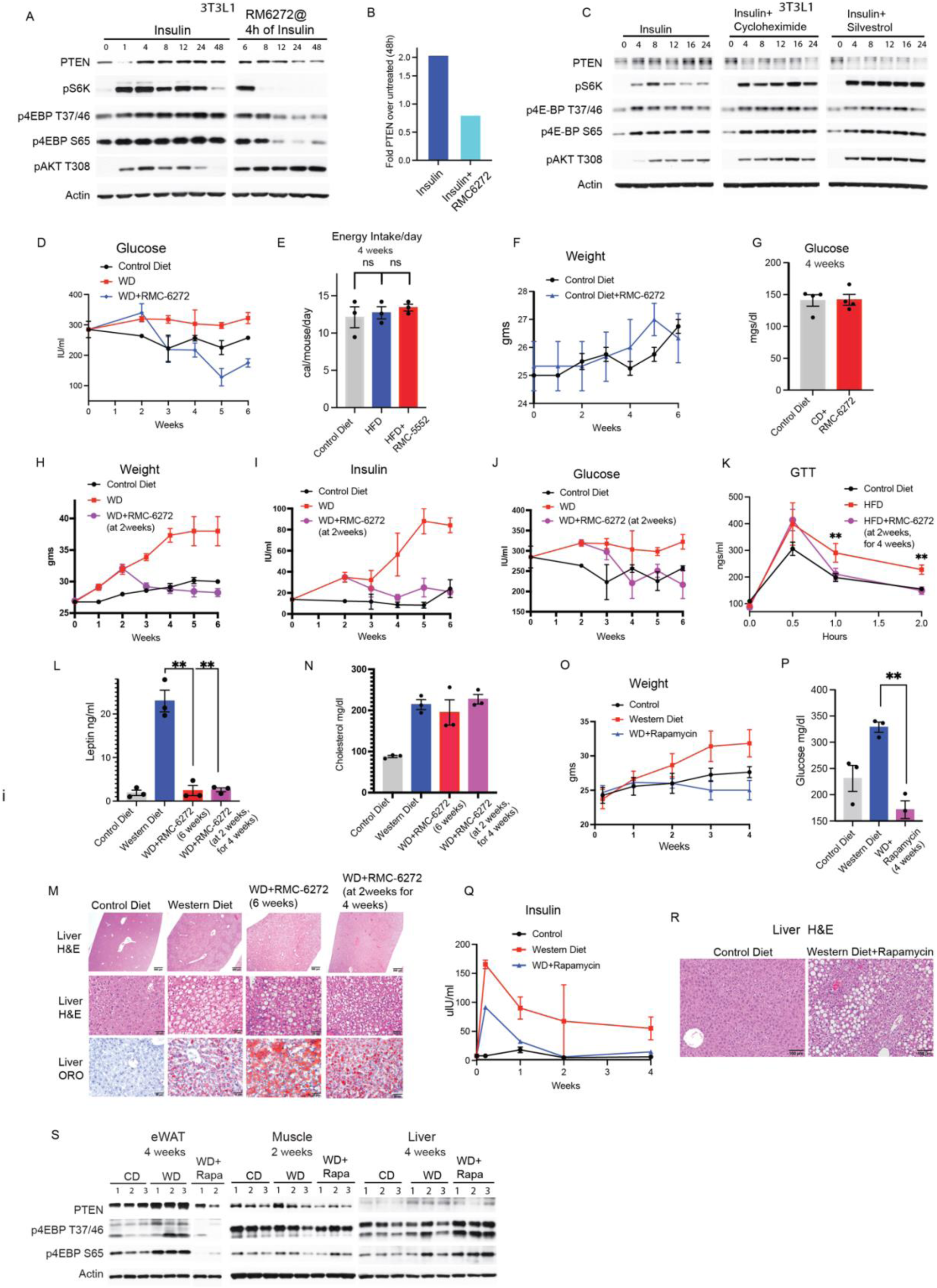
PTEN increase is driven by mTORC1 dependent translation and inhibiting mTORC1 prevents and reverses insulin resistance. **(A)** 3T3L-1 cells were treated insulin alone or with RMC-6272 (0.5nM), 4 hours after insulin for the indicated times and immunoblotted for PTEN and PI3K pathway activity. **(B)** Quantification of PTEN from A and represented as fold change over untreated control. **(C)** 3T3L1 cells were treated with1ug/ml of insulin alone or with a combination of insulin and cycloheximide (30ug/ml) or silvestrol (30nM) and immunoblotted for PTEN and PI3K pathway activity. **(D)** Mice were fed on regular diet or western diet or western diet treated with the mTORC1 inhibitor RMC-6272 (3mg/kg) once a week for 6 weeks and the glucose levels p< 0.01(n=5) measured. **(E-G)** Mean and SEM of energy intake, weight and glucose measured of mice for the indicated times and conditions (n=3). **(H-K)** Mice were fed on the indicated diets for 2 weeks and then treated with RMC-6272 (3mg/kg) on western diet for 4 weeks, weight p<0.0001 (n=5-15), insulin p<0.0005 (n=5), glucose levels (n=5) and GTT measured. **(L, N)** Leptin and cholesterol was measured for the indicated times and treatment conditions. **(M)** Livers from mice with indicated diet and treatment stained for H&E and lipid OilRedO (n=3). Two independent experiments done and representative images are shown. **(O-Q)** Weight (p<0.0001), glucose and insulin (p<0.01) was measured in mice fed with control diet, western diet or western diet with Rapamycin treatment (10mg/kg) for the indicated times (n=3). **(R)** Livers from mice fed with control diet or western diet with rapamycin treatment were stained for H&E (n=3). Two independent experiments done, and representative images are displayed. **(S)** eWAT, muscle and liver of mice fed and treated with indicated diets and rapamycin analyzed by immunoblotting for PTEN levels and mTORC1 activity. Mean and SEM of n plotted, p values calculated by one-way ANOVA test (K), by two-way ANOVA and post Bonferroni tests (D, H-I, O, Q), by student’s t test (J, L, N, P). *p<0.05, **p<0.01, ***p<0.001, ****p<0.0001.

